# CAM: a quality control pipeline for MNase-seq data

**DOI:** 10.1101/143263

**Authors:** Sheng’en Hu, Xiaolan Chen, Ji Liao, Yiqing Chen, Chengchen Zhao, Yong Zhang

## Abstract

Nucleosome organization affects the accessibility of cis-elements to trans-acting factors. Micrococcal nuclease digestion followed by high-throughput sequencing (MNase-seq) is the most popular technology used to profile nucleosome organization on a genome-wide scale. Evaluating the data quality of MNase-seq data remains challenging, especially in mammalian. There is a strong need for a convenient and comprehensive approach to obtain dedicated quality control (QC) for MNase-seq data analysis. Here we developed CAM, which is a comprehensive QC pipeline for MNase-seq data. The CAM pipeline provides multiple informative QC measurements and nucleosome organization profiles on different potentially functional regions for given MNase-seq data. CAM also includes 268 historical MNase-seq datasets from human and mouse as a reference atlas for unbiased assessment. CAM is freely available at: http://www.tongji.edu.cn/~zhanglab/CAM

## Introduction

Nucleosome organization (i.e., the relative location of a nucleosome on the DNA) affects the transcriptional activity by influencing the access of DNA-binding proteins to the genome and the elongation of RNA polymerase II [1, 2]. Recently, nucleosome organizations in a variety of species and cell types have been profiled using micrococcal nuclease digestion followed by high-throughput sequencing (MNase-seq) [3]. Although MNase-seq technology has been widely used and many computational tools have been developed for MNase-seq data [4], the quality evaluation remains challenging, especially for data from mammalian genomes, due to two major difficulties. First, different experimental designs (e.g., sequencing coverage and the MNase concentration) may result in distinct nucleosome organization features in some genomic loci (e.g., fragile nucleosomes at promoters [5]). Second, in contrast to chromatin immunoprecipitation sequencing (ChIP-seq), DNase-seq and methylated DNA immunoprecipitation sequencing (MeDIP-seq) data, the MNase-seq data signals are not enriched in any specific genomic loci, resulting in difficulties in focusing on target regions for downstream analysis. Many software tools were designed to detect well-positioned nucleosomes, but seldom took care of MNase specific quality control which is the basis for detecting nucleosome organization correctly and precisely. Here, we present CAM, an integrated quality control (QC) for MNase-seq data. CAM provides multiple key measurements that enable users to evaluate the data quality using scores from 268 historical MNase-seq datasets in human and mouse as a reference atlas. In addition, CAM provides nucleosome organization information based on potentially functionally related genomic regions for use in the targeted downstream analysis.

## Results and conclusion

### Overview of CAM

The CAM pipeline initiates from the data pre-processing steps, including reads mapping (optional), high-quality reads filtering (optional) and nucleosome organization profile generation (Fig 1). After the pre-processing steps, CAM provides multiple QC measurements to allow users to evaluate the data quality as follows: 1) sequencing coverage, 2) AA/TT/AT dinucleotide frequency, 3) nucleosomal DNA length, 4) existence of nucleosome free regions (NFR) at promoters, 5) well-positioned nucleosomes at the downstream promoters, 6) well-positioned nucleosomes at custom defined potential cis-regulatory regions, 7) enrichment of well-positioned nucleosome arrays in DNase hypersensitive sites (DHS) (S1 Table). We compiled 268 MNase-seq datasets from human and mouse as a historical QC reference atlas and made an unbiased judgment for the measurements by generating several QC scores (Method section, S2 Table). In addition, CAM provides related analysis results including 1) nucleosome profiles at each of promoter and custom defined region, 2) detection of well-positioned nucleosome arrays, 3) gene level annotation of the well-positioned nucleosome arrays.

**Fig. 1.**
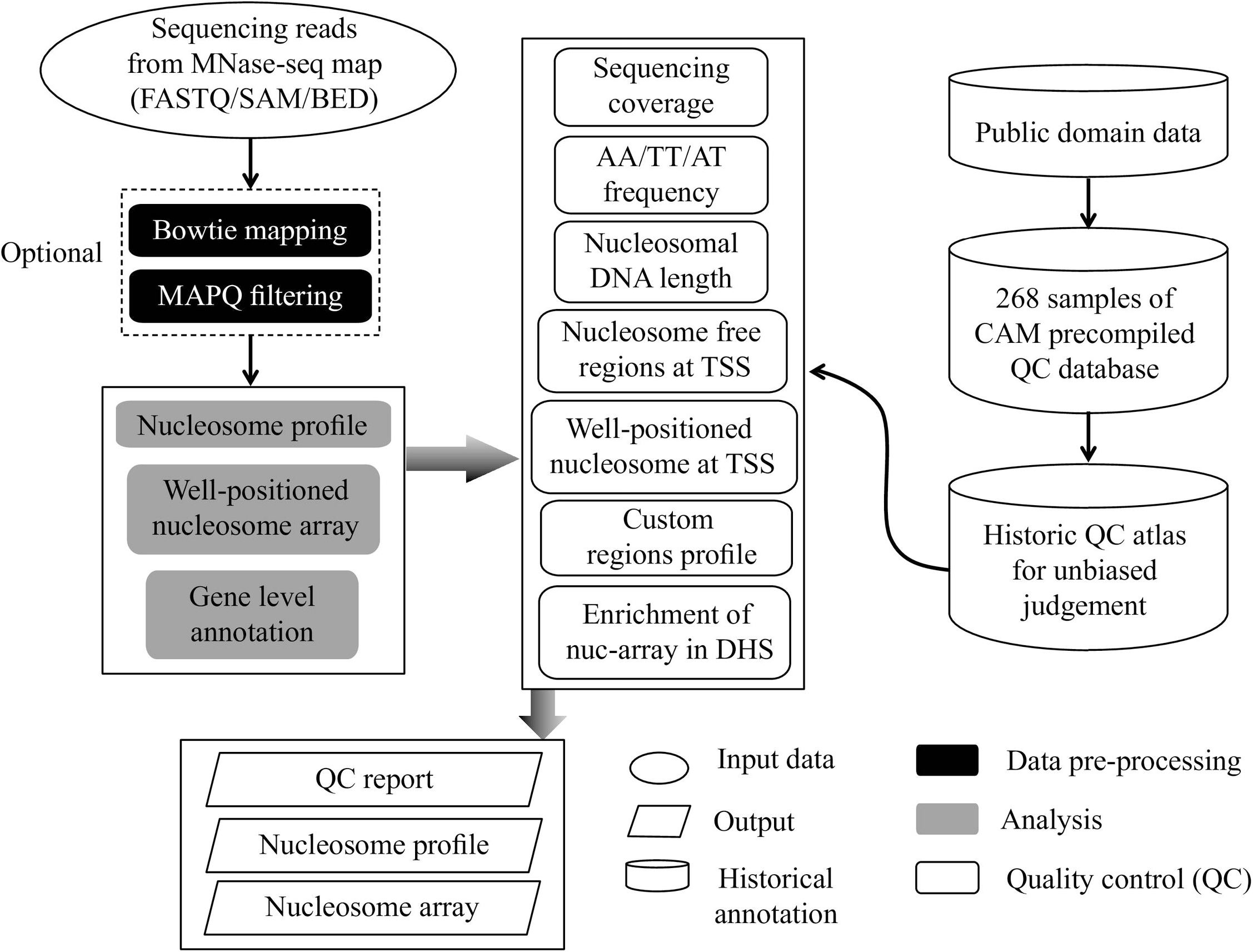
Flowchart illustrating the CAM with default parameters. The workflow of CAM includes three components: data pre-processing (black box), analysis (grey box) and QC (white box). The QC metrics for all historical MNase-seq data were precompiled for unbiased judgment in the QC component.

Many software tools have been developed for MNase-seq data analysis. However, most of them focused on detecting positioned nucleosomes and did not take care of the quality of MNase-seq data, which has great importance on nucleosome detection. Additionally, compared with the concept of positioned nucleosome, array of well-positioned nucleosome provides a more unbiased description of nucleosome positioning detection for MNase-seq data [1, 6–8] (discussed in the following section) (S3 Table). CAM has three advantages over existing state-of-the-art methods: 1) CAM provides QC components specific to MNase-seq. 2) Like most of existing software tools, CAM also provides a peak calling function, however the peak calling function is based on a more unbiased concept: well-positioned nucleosome array. 3) CAM is implemented as a systematic pipeline, which takes raw MNase-seq reads or aligned reads as input, and outputs QC measurements together with a series of analysis results (discussed in the following section).

## Historical QC reference atlas from published MNase-seq data

To provide an unbiased judgment of QC measurements, we collected 268 MNase-seq samples from human and mouse and pre-processed with all of our QC components. Sequencing coverage, rotational score, estimated fragment length and nucleosome profiles at promoters were collected as historical QC reference to measure the quality of input MNase-seq data (S2 Table).

## Sequencing coverage measures the resolution of nucleosome positioning

Sequencing coverage provides a direct measurement of the resolution of two features of nucleosome organization (i.e., occupancy and positioning) [9]. We generated simulated regions with perfect positioning and no positioning, and compared the difference of nucleosome positioning scores with different sequencing coverage to show the influence of sequencing coverage on the resolution of nucleosome positioning detection (Method section). The perfect positioning region and no positioning region showed similar positioning scores with low sequencing coverage (almost the same between them in 1-fold coverage) and the nucleosome positioning score of the perfect positioning region significantly increased compared with the no positioning region with higher sequencing coverage (Fig 2A). In the CAM pipeline, we calculated sequencing coverage of the input MNase-seq data and compared it with the distribution of historical data (Fig 2B) to reflect the ability of MNase-seq data to detect nucleosome positioning and occupancy.

**Fig. 2.**
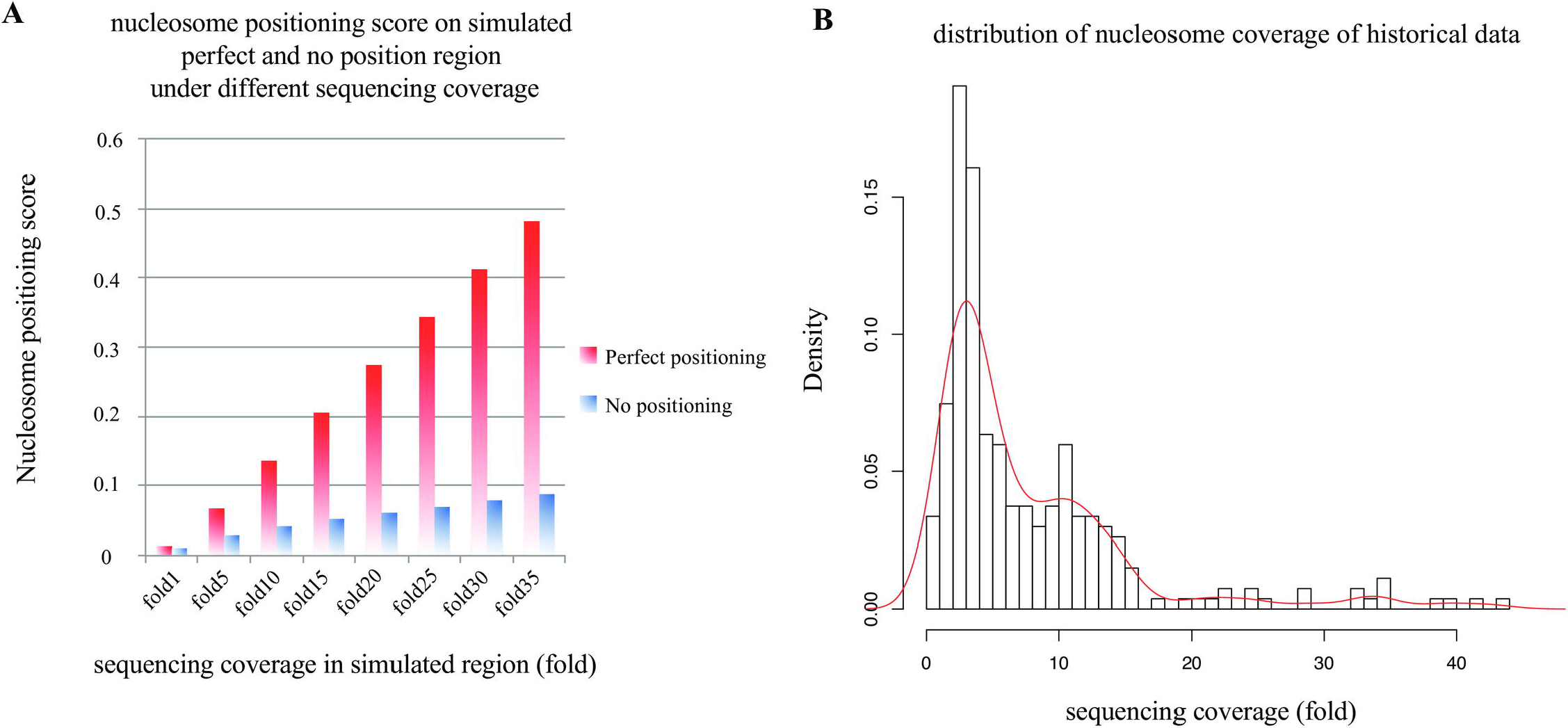
Average nucleosome coverage. **(A)** Regions with higher sequencing coverage exhibit higher resolution for the detection of nucleosome positioning. The barplot shows the nucleosome positioning score in a simulated perfect positioning region and no positioning region. For low sequencing coverage, the positioning score is almost the same for the perfect and no positioning regions, and the difference increases when the coverage increases. **(B)** Distribution of sequencing coverage of all historical data.

## AA/TT/AT periodicity measures the nucleosome rotational positioning

The 10-base AA/TT/AT di-nucleoide periodicity in the nucleosomal DNA provides a measurement of nucleosome rotational positioning, which is influenced by the DNA sequence. Studies show that AA/TT/AT di-nucleotide displays a 10 bp periodicity throughout nucleosomal DNA sequence. The di-nucleotide favors DNA bending in a specific direction and expands the major groove [10]. Thus the 10 bp periodicity should be observed in the AA/TT/AT di-nucleotide frequency of MNase-seq reads if the MNase-seq reads reflect nucleosomal DNA accurately. We defined a “rotational score” to measure the 10 bp AA/TT/AT periodicity (Method section) and compared with the distribution of historical data (Fig 3A). A judgment of “rotational score” (Pass or Fail) was assigned for the input sample to measure the rotational positioning (Fig 3B).

**Fig. 3.**
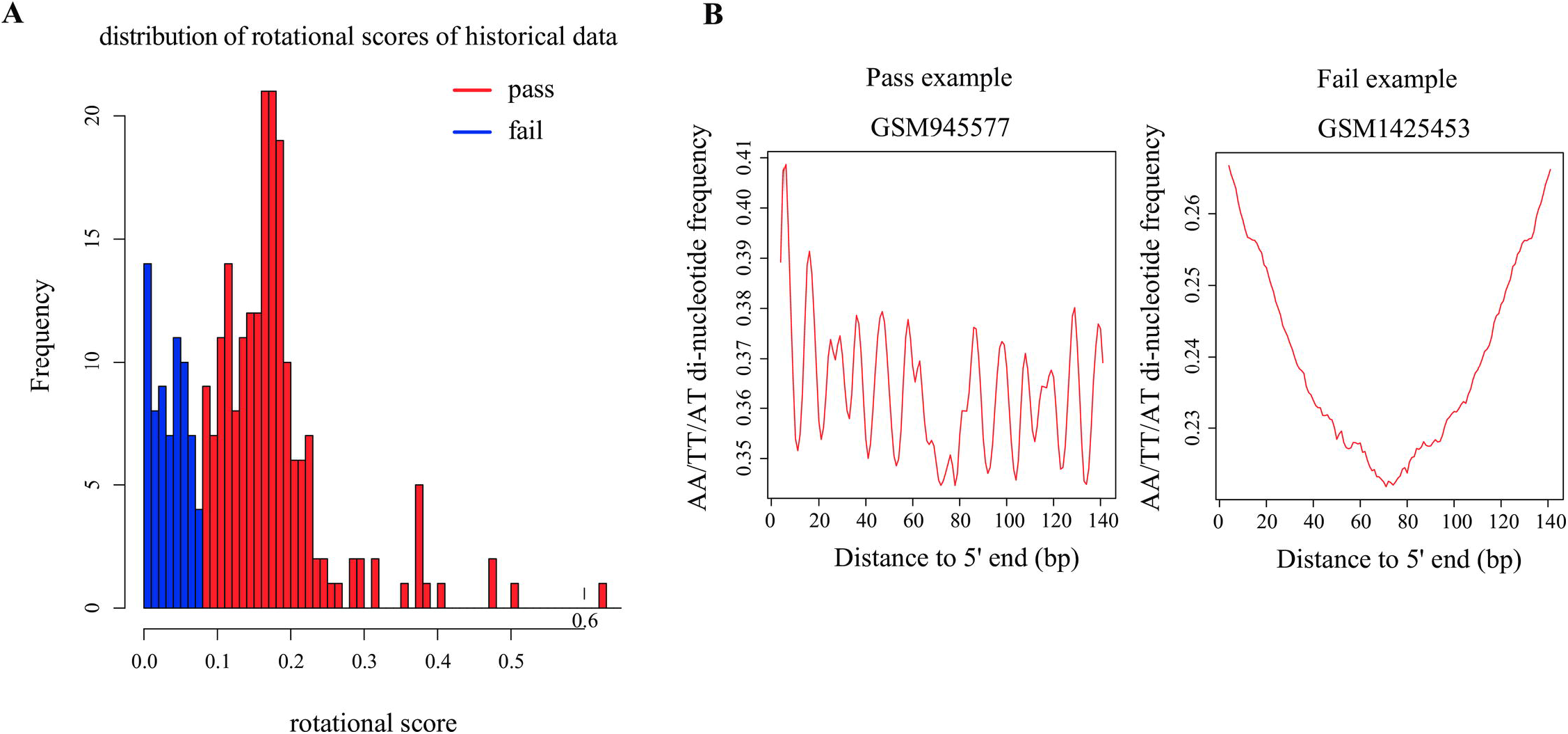
AA/TT/AT di-nucleotide periodicity. **(A)** Distribution of the rotational scores of all historical data. A rotational score < 0.08 was determined as a “Fail” (blue) in this measurement. **(B)** Examples of “Pass” and “Fail” MNase-seq data in this measurement.

## The nucleosomal DNA length distribution reflects the MNase concentration

The nucleosomal DNA length distribution (referring to the fragment length or MNase library size) is closely related and thus reflects the MNase concentration. MNase concentration is negatively correlated with the fragment length estimated from the MNase-seq data: Samples with higher MNase concentration exhibited shorter nucleosomal DNA length, whereas lower concentration samples exhibited longer nucleosomal DNA length (Fig 4A). CAM estimates fragment length of nucleosomal DNA from the input data and compared with the distribution of historical data to reflect the MNase concentration (Method section, Figure 4B). Although different MNase concentration may result in different nucleosome organization features in some genomic loci (e.g. fragile nucleosomes at promoters) [5], we expect the fragment length to be close to the length of nucleosomal DNA (147 bp), indicating that the MNase-seq reads accurately represent the location of mono-nucleosome (through a judgment of Pass or Fail, Fig 4B, C). A longer or shorter fragment length indicates partial or over digestion of MNase, respectively.

**Fig. 4.**
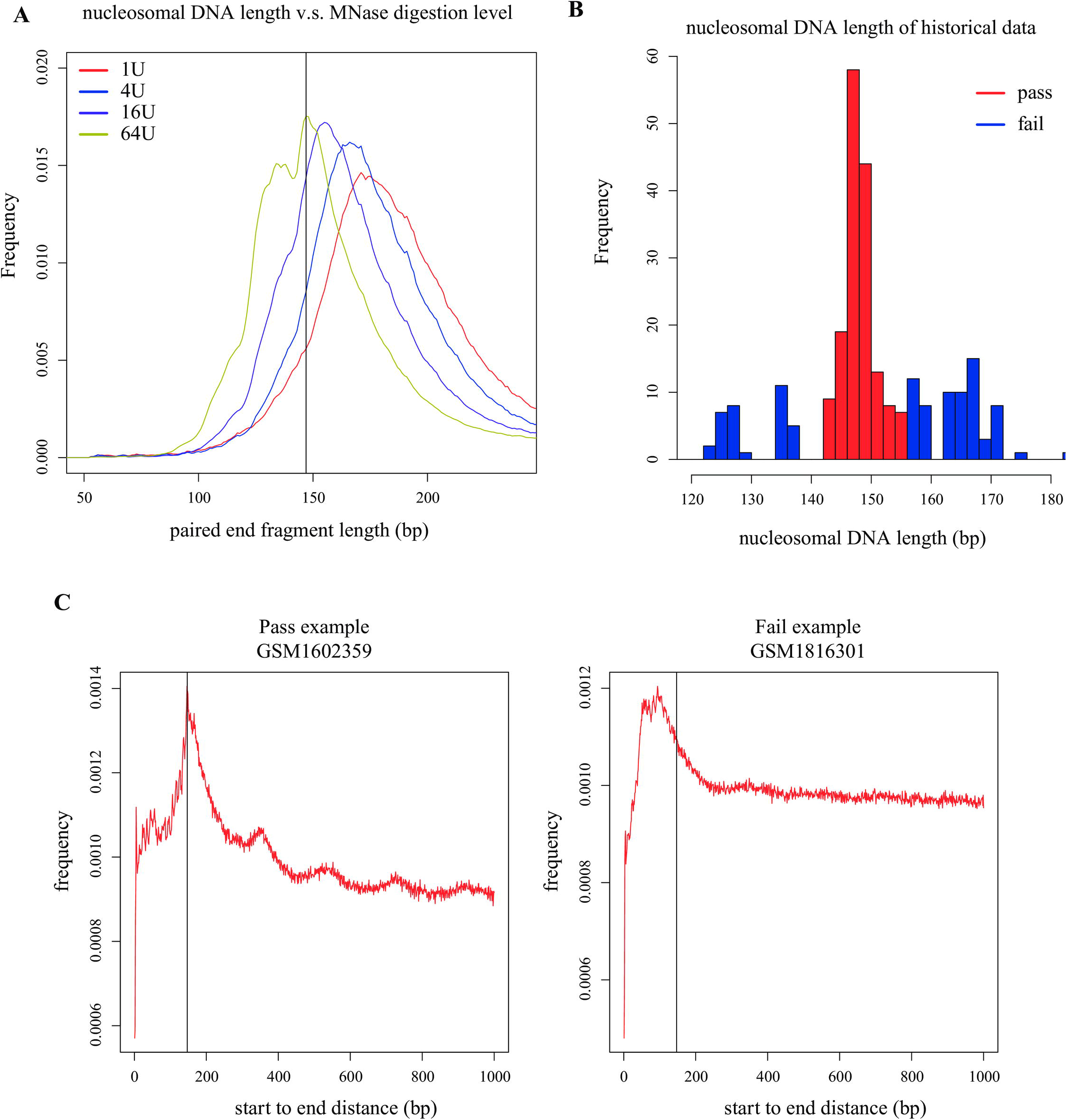
Nucleosomal DNA length distribution. **(A)** The MNase digestion level (concentration) is related to the nucleosomal DNA length: samples with a higher MNase concentration exhibite shorter nucleosomal DNA length, whereas lower concentration samples exhibite longer nucleosomal DNA length. The MNase-seq data used for the comparison were obtained from a previous study in mouse ESCs (GSM2083105, GSM2083106, GSM1083107, GSM1083108). **(B)** The distribution of the nucleosome length from all of the historical data. Nucleosome length < 140 or > 155 was determined as “Fail” (blue) in this measurement. **(C)** Examples of MNase-seq data with “Pass” and “Fail” in this measurement. The vertical line labels 147 bp.

## Nucleosome profiles on potentially functional regions reflect the ability for detecting well-positioned nucleosomes

Well-positioned nucleosome, an effective marker of transcriptional regulatory regions, is formed by nucleosomes consistently positioned in a cell population. Thus, MNase-seq reads can be observed to position consistently in certain functional regions to form well-positioned nucleosomes. Promoter regions are regarded as the most important functional regions for transcriptional regulation. According to the barrier nucleosome model [1, 2], nucleosome free regions (NFR) and the successive well-positioned nucleosomes are supposed to be observed around TSS. CAM generates nucleosome profiles on promoter regions and displays with an aggregate plot and a heatmap (Fig 5A). Additionally, CAM calculates two QC scores according to the nucleosome profiles on promoter regions, and assigns judgments for the input sample to measure the nucleosome positioning pattern at promoter regions, based on the comparison with the distribution of the historical data (Method section). The two QC scores consist of 1) a promoter NFR score to check the existence of NFR at promoters (Fig 6A, B) and 2) a promoter positioning score to measure the successive well-positioned nucleosomes at the downstream promoters (Fig 6C, D).

**Fig. 5.**
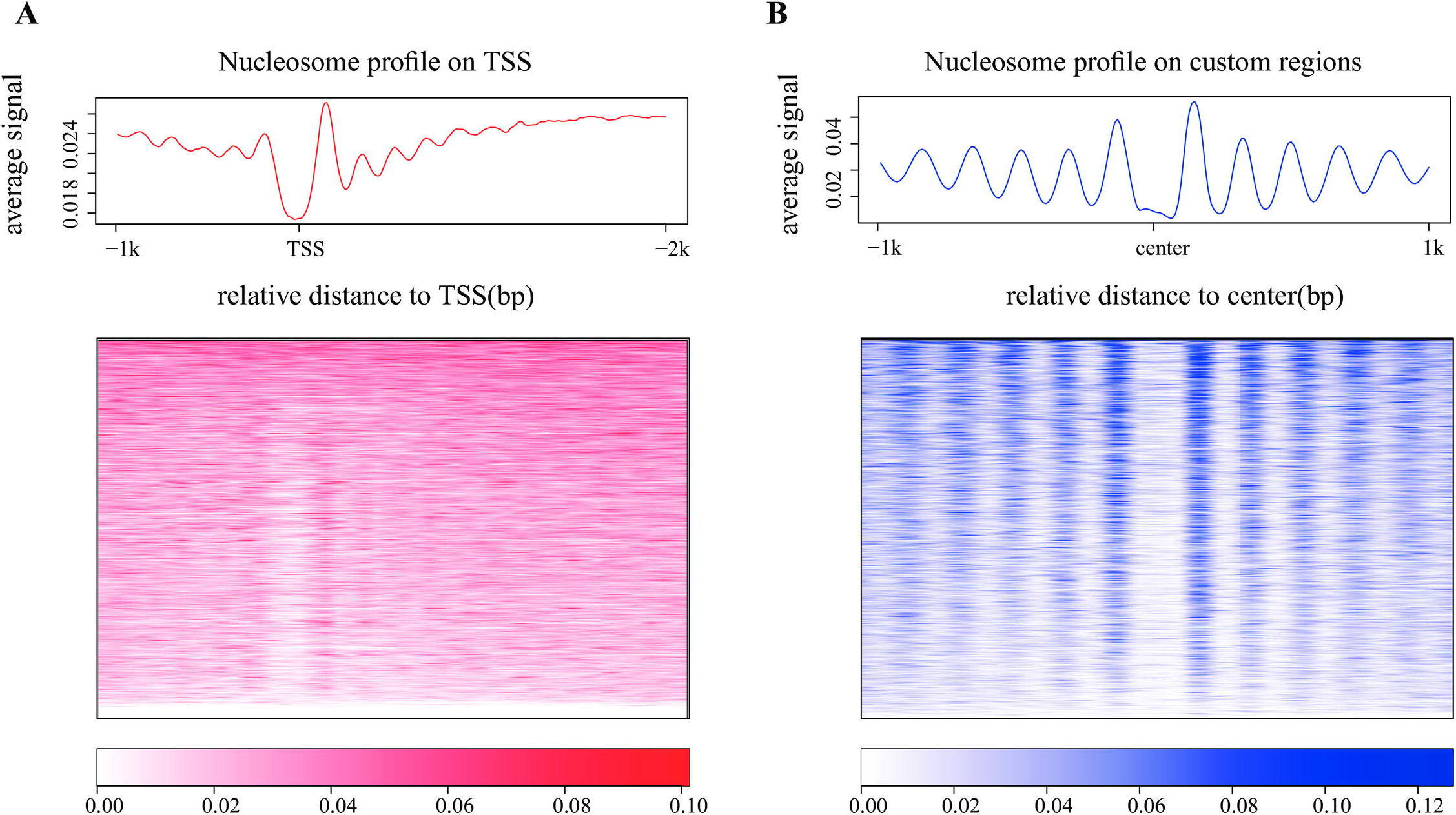
Nucleosome profiles on promoter regions and custom regions. Nucleosome profiles were generated and plotted as aggregate plots and heatmaps on **(A)** promoters and **(B)** custom regions. By default, the promoter regions were defined as -1 kb to +2 kb from the TSS of all refseq genes, and the custom regions were extended to +/-1 kb (by default) from the center of the regions. Both the promoters and custom regions were sorted by the MNase-seq read counts within the regions. An MNase-seq data from a human lymphoblastoid cell line (GSM907784) was selected to plot the nucleosome profiles on both promoters and custom regions.

**Fig. 6.**
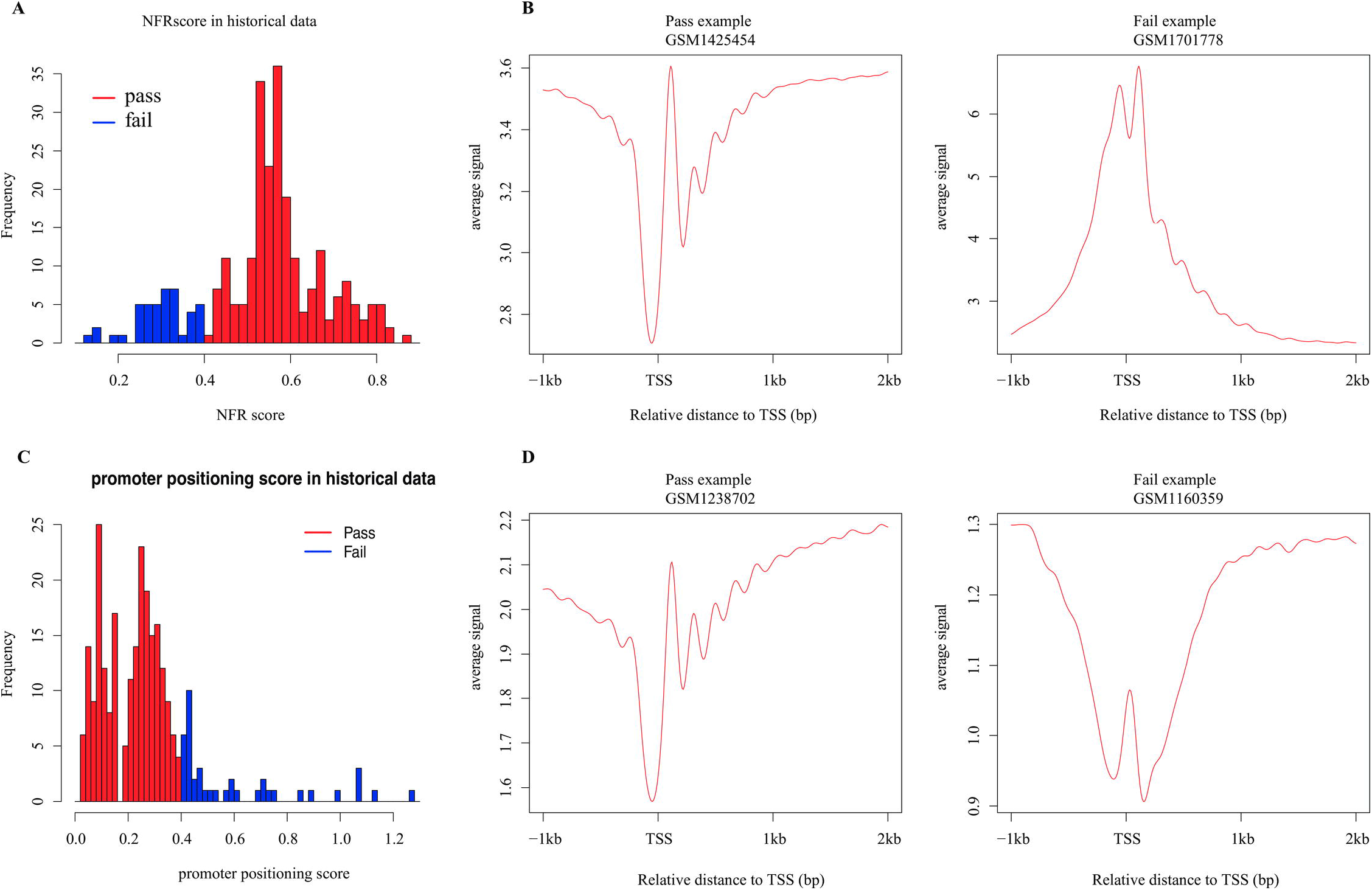
Distribution of promoter NFR score and promoter positioning score. Distribution of the promoter NFR score of all historical data. A promoter NFR score < 0.4 was determined as a “Fail” (blue) in this measurement. (B) Examples of “Pass” and “Fail” MNase-seq data in promoter NFR score. (C) Distribution of the promoter positioning scores of all historical data. A promoter positioning score > 0.4 was determined as a “Fail” (blue) in this measurement. (D) Examples of “Pass” and “Fail” MNase-seq data in promoter positioning score.

In addition, users can provide custom regions for CAM to profile nucleosome organization at the same time. For example, users can provide motif sites or binding sites (defined by ChIP-Seq peaks) of certain transcription factors (CTCF motif sites as example, Fig 5B) as custom regions, and check whether the successive well-positioned nucleosomes are also observed at these regions as an additional QC measurement.

In our previous work, we developed an effective method to detect cis-regulatory regions with MNase-seq data, which we called the detection of well-positioned nucleosome arrays [8]. A well-positioned nucleosome array is a broader region with successive well-positioned nucleosomes, often emanate from a nucleosome-depleted region, such as transcription factor binding site [1, 6, 7], and is very unlikely to be generated at random.

CAM adopts the method to detect nucleosome arrays across whole genome (Method section, Fig 7A displays an example of detected well-positioned nucleosome arrays) [8]. The nucleosome arrays define a list of potential target regions for the following analysis, which is similar as ChIP-Seq peaks, and solve the problem that MNase-seq data do not show enrichment in any regions. We showed that these nucleosome arrays were enriched in potentially functional regions such as the downstream promoters and the union DHS sites (Method section, Fig 7B), which can be explained by the barrier nucleosome model [11]. Thus we calculated the fold enrichment of the well-positioned nucleosome arrays on the union DHS sites compared with random background (Method section) as a QC measurement to assess the ability of the MNase-seq data to detect cis-regulatory elements with well-positioned nucleosome arrays. Samples with fold enrichment less than 2 are regarded as “Fail” in this measurement, indicating the well-positioned nucleosome arrays are more likely to be caused by random rather than the barrier nucleosome model. The genomic coordinates together with the nucleosome profile values are reported for each region with a well-positioned nucleosome array. CAM also provides a gene level annotation of the well-positioned nucleosome arrays for downstream analysis (Method section).

**Fig. 7.**
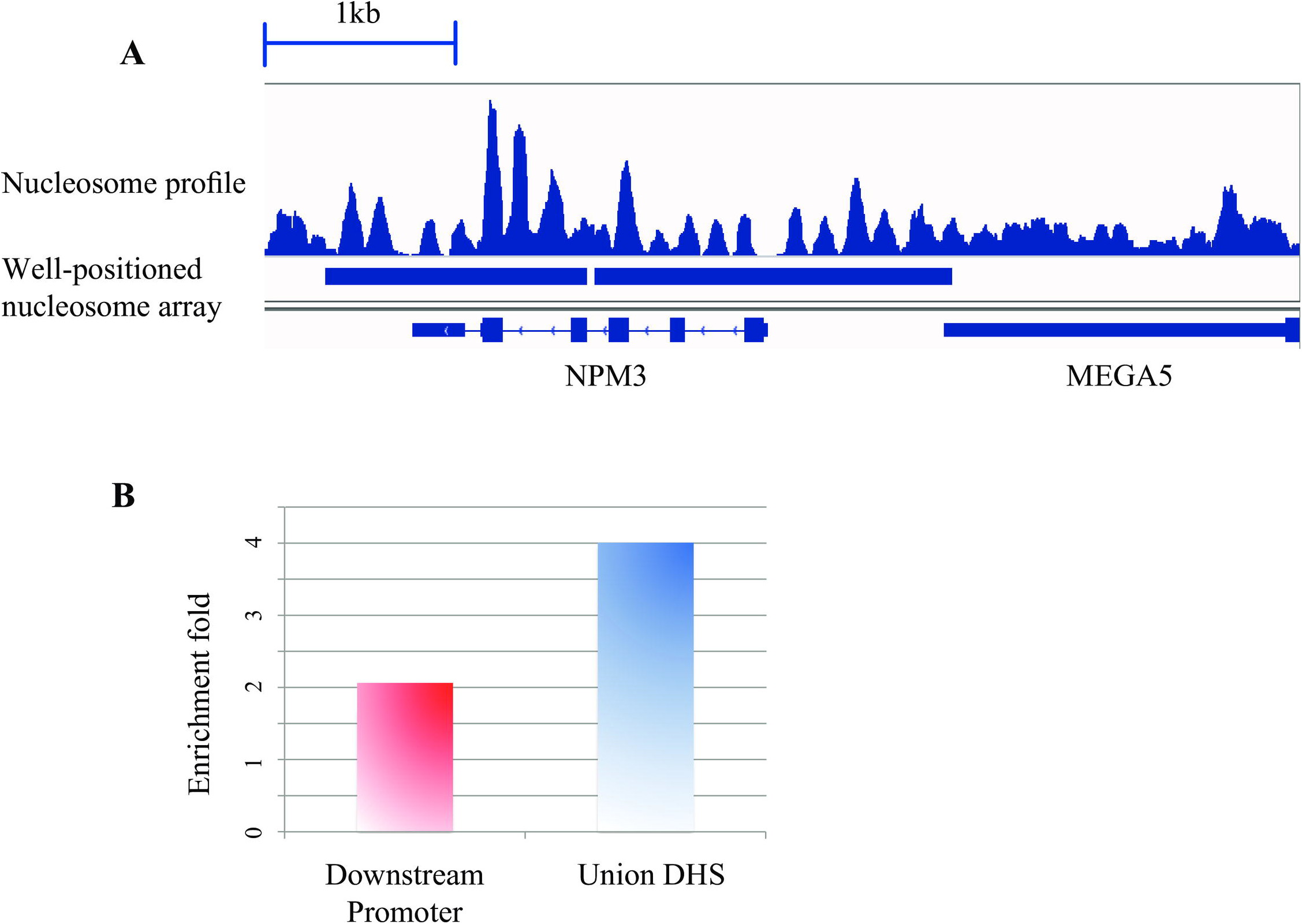
Well-positioned nucleosome array. **(A)** An example of detected well-positioned nucleosome arrays. An MNase-seq data from a human lymphoblastoid cell line (GSM907784) was selected to plot the nucleosome profile for the example region. **(B)** Well-positioned nucleosome arrays are enriched in both the downstream promoters and the union DHS sites. The enrichment fold (y-axis) represents the fold change of the observed proportion of nucleosome arrays on the downstream promoters (or the union DHS sites) to the expected proportion (Method section). The nucleosome arrays were detected using MNase-seq data from human lymphoblastoid cell line (GSM907784).

## Computational cost and standard output of CAM

We applied CAM to published MNase-seq data from a human lymphoblastoid cell line (GSM907784) and obtained multifaceted and detailed QC reports (S1 File) together with a series of analysis results (S4 Table). Running time the CAM pipeline is also provided (S5 Table).

## Conclusion

In summary, CAM is specifically designed for QC of MNase-seq data. The QC components measure the quality of MNase-seq data in different aspects (S1 Table). The program uses standard format input files via simple commands, reports informative QC measurements (specific for MNase-seq data) to assist in evaluating the data quality (using historical MNase-seq data as a reference atlas), generates nucleosome organization profiles based on promoters and custom defined regions, and detects regions with well-positioned nucleosome arrays.

## Materials and Methods

### Implementation and webpages of CAM

CAM was implemented using Python and R. Users need python (version = 2.7) and R (version >= 2.14.1) installed on linux or MacOS environment to make sure the successful process of CAM. CAM was distributed under the GNU General Public License. The source code and detailed tutorial of CAM is available on our webpages: http://www.tongji.edu.cn/~zhanglab/CAM.

### Data pre-processing

In the alignment process, Bowtie (Langmead et al., 2009) was used to align the MNase-seq reads with the −m 1 parameter. Users could adjust two other parameters (“-X” for maximum insert size in paired end data and “-3” for trimming bases from 3’end of reads) in the configure file. To maintain high-quality alignment results, we removed reads with a sequencing quality less than 30 (The default cutoff of MAPQ filtering is 30). The alignment step was skipped when the aligned reads (SAM/BED format) input was used. CAM was designed specifically for MNase-seq data from human and mouse and only supports (by default) the hg38, hg19, mm10, and mm9 genome versions. Users can add custom genome versions according to the instructions in the CAM manual.

To generate nucleosome profiles, the sequencing reads were transformed into nucleosome reads as follows: for single end sequencing data, the reads were extended to 147 bp in the 3’ direction; for paired end data, the paired end fragments were extended to 73 bp in both the 5’ and 3’ directions from the fragment centers. Then, the middle 73 bp centered on the extended fragments were compiled as the nucleosome profile.

### Calculation of sequencing coverage

The sequencing coverage (fold) was defined as (*Number of reads* × 194 *bp*)/(*Effective genome size*). The “number of reads” was defined as the number of mappable reads after MAPQ filtering for single end data and the number of fragments for paired end data. Additionally, “194 bp” represented the total length of the nucleosomal DNA and linker DNA, which was estimated from 268 historical MNase-seq datasets. The “effective genome size” was defined as 2.7 billion nucleotides (2.7e9) for human and 1.87 billion nucleotides (1.87e9) for mouse. The effective genome size used here was smaller than the original genome size due to the repetitive features on the chromosomes.

### Simulation of the perfect positioning and no positioning regions

We compared the nucleosome positioning scores on a simulated perfect positioning region with a no positioning region with different sequencing coverage to demonstrate the influence of the sequencing coverage on the resolution of nucleosome positioning. A 1200-bp region was prepared for all simulations. Five “potential nucleosome centers” were marked with a distance between adjacent centers equal to 194 bp (nucleosomal + linker DNA length, estimated with the historical data). Simulated reads (1 bp read to reflect the nucleosome center) were assigned evenly to each of the 5 “potential nucleosome centers” in the perfect positioning region with a 5 bp perturbation. For the no positioning region, simulated reads were assigned randomly on the 1200 bp region with equal probability and the same perturbation as the perfect positioning region (5 bp). The nucleosome positioning scores for the different sequencing coverage in the two types of regions were calculated using the method described below (“Detect well-positioned nucleosome arrays” section).

### Measurement of AA/TT/AT di-nucleotide frequency and definition of rotational score

We sampled down all mappable reads to 10 million and extended each read to 147 bp in the 3’ direction. Then, the aggregate AA/TT/AT di-nucleotide frequency was calculated across 4 to 143 bp of the extended reads. We conducted a Fourier transform on the aggregate frequency and used the energy of 10-bp periodicity (defined as the rotational score) to demonstrate the extent to which the MNase-seq reads reflect the nucleosome organization. Samples with rotational scores greater than 0.08 were defined as “Pass” in this measurement, whereas the other samples were defined as “Fail”. The cutoff 0.08 was determined from the distribution of rotational scores among all historical MNase-seq data.

### Calculation of nucleosomal DNA length distribution

For the paired end samples, the fragment length distribution from all mappable fragments was used to directly infer the nucleosomal DNA length distribution. For the single end samples, we calculated a start-to-end distance to estimate the nucleosomal DNA length as follows: mappable reads were sampled down to 10 million; then, we calculated the distribution of the distance from the 5’ end of each plus strand read to all 5’ ends of the minus strand reads (the start-to-end distance) within 250 bp downstream (1kb downstream in figures for visualization). Duplicate reads were discarded in this calculation. After the distribution of the start-to-end distance was generated, the length with the highest frequency was defined as the estimated nucleosomal DNA length of the MNase-seq data (for both the paired end and single end data). Samples with estimated nucleosomal DNA lengths in the range of 140 to 155 bp were defined as “Pass”, whereas the other samples were defined as “Fail”. The cutoff was determined from the distribution of the nucleosomal DNA lengths among all historical MNase-seq data.

### Nucleosome organizations on the promoters and custom regions

The nucleosome profile was generated as described in the “Data pre-processing” section. The nucleosome signal from 1 kb upstream to 2 kb downstream (default, user can change the range with certain parameters in the configure file; see Manual for details) from TSS was plotted as a heatmap and aggregate curve with a 10-bp resolution. For custom regions, the nucleosome signal was plotted within +/− 1kb (default) from the center of the region. The signals from the minus strand transcripts and the minus strand custom regions were reversed in both the heatmap and the aggregate curve.

### Calculation of promoter NFR score and promoter positioning score

To calculate the promoter NFR score, we first generated the average MNase-seq signal profile from 1kb upstream to 2kb downstream from all TSS. The aggregate signal was then kernel smoothed to remove signal noise. Promoter NFR score was calculated by the smoothed aggregate signal of the +1 nucleosome and the -1 nucleosome subtracting the signal of nucleosome free region. Finally, the promoter NFR score was normalized by the difference between the maximum signal and the minimum signal within the 3kb promoter regions. We regarded aggregate signal of the first downstream local maximum as the signal of +1 nucleosome, the first upstream local maximum as -1 nucleosome and the first upstream local minimum as NFR.

The promoter positioning score was defined as the coefficient of variance (CV) of the distance between +1, +2, +3 and +4 nucleosomes. The position of nucleosomes was defined as the local maximum positions.

### Detection of well-positioned nucleosome arrays

To detect well-positioned nucleosome arrays, first, the mappable read were extended to 147 bp in the 3’ direction; then, the centers of the extended reads were compiled to generate the nucleosome center profile (for the paired end data the center of each fragment was compiled directly). Next, Gaussian smoothing (window size = +/-73 bp; standard deviation = 30) was performed on the nucleosome center profile, and the absolute difference between the adjacent bps was calculated as the modified profile. Then, the local maximum within +/-73 bp was selected. All adjacent local maxima were connected to create a “nuc-array” curve and a signal that was defined as the “nuc-array” value. The “nuc-array” value was transformed to a fold enrichment value over the background (defined as the average “nuc-array” value across the genome). The well-positioned nucleosome array was defined as segments with lengths greater than 600 bp and fold enrichment greater than 2 (default, the cutoff of length and fold enrichment can be changed by the users). We generated a 5 columns bed file with the above method (named as “outputname_Nucleosome_Array.bed”). Each line of the bed file represented a well-positioned nucleosome array. The 5th column of the bed file is the “nuc-array” value. The method was also described in a previous work [8]. The positioning score for the simulated regions (described above) was calculated by the average “nuc-array” value across the whole simulated regions.

### Enrichment of well-positioned nucleosome arrays in downstream promoter regions and union DHS sites

The enrichment fold on the promoters of the nucleosome array was defined as the fold change of observed and expected percentage of the well-positioned nucleosome arrays on promoters. The expected percentage was equal to the percentage of the promoter length compared with the effective genome size (mentioned above). Then, the promoter regions were defined as 3 kb downstream from the TSS of refseq genes. Similar enrichment fold was also performed on the Union DHS sites, which were generated by merging the narrow peak of all DNase-seq data from the human and mouse (separately) from the ENCODE and Roadmap Epigenomics project as previously described [12]. Samples with enrichment fold greater than 2 were defined as “Pass” in this measurement, otherwise they were “Fail”.

### Gene level annotation of well-positioned nucleosome arrays

Well-positioned nucleosome arrays are annotated to genes by overlapping with the promoters of genes, which are defined as 3 kb upstream and downstream from TSS. The length of overlapped nucleosome arrays is marked as a feature of the certain gene (“promoter nuc-array length”). CAM provides an additional analysis result in the output folder, which is named as “outputname_geneLevel_nucarrayAnnotation.bed”. The additional analysis result is a bed file with 6 columns, including “chromosome name”, “promoter start site”, “promoter end site”, “refseq ID”, “promoter nuc-array length” and “strand”. Each row of the bed file represents a refseq transcript. The 5th column (“promoter nuc-array length”) represents the length of the well-positioned nucleosome array which overlaps with the certain gene promoter (-1 for no nucleosome array overlapped). With the help of this analysis result from different samples, users can easily compare the array length of same genes in different conditions.

## Acknowledgements

We thank Kai Fu and Jiangxing Feng for their contributions in the early stage of this project.

## Supporting Information

**S1 File. CAM QC reports with default parameter for MNase-seq data from human lymphoblastoid cell line (GSM907784).**

**S1 Table. Summary of QC measurements in CAM.**

**S2 Table. Meta data, QC scores and accession ID for 268 samples of historical MNase-seq data.** Sequencing coverage, rotational scores, estimated fragment length, promoter NFR scores and promoter positioning scores. The scores and judgments for historical MNase-seq data were also attached.

**S3 Table. Compare of function between CAM and other nucleosome analysis software.** We compare the major function of CAM to existing state-of-the-art methods. CAM has three advantages: 1) Quality control component specific to MNase-seq, 2) Well-positioned nucleosome arrays detection and 3) Systematic pipeline for users to input raw sequencing data and get all QC and analysis results.

**S4 Table. List of standard output from CAM.**

**S5 Table. Running time of CAM.** An MNase-seq data from human lymphoblastoid cell line (GSM907784, totally 546,924,994 reads) was used to evaluate the runtime of CAM. Alignment process was excluded from this calculation. The percentage running time for each component was calculated by using single CPU (Intel^®^ Xeon^®^ CPU E5-2640 v2 @ 2.00 GHz).

